# Design Specifications for Cellular Regulation

**DOI:** 10.1101/081729

**Authors:** David C. Krakauer, Lydia Müller, Sonja J. Prohaska, Peter F. Stadler

## Abstract

A critical feature of all cellular processes is the ability to control the rate of gene or protein expression and metabolic flux in changing environments through regulatory feedback. We review the many ways that regulation is represented through causal, logical and dynamical components. Formalizing the nature of these components promotes effective comparison among distinct regulatory networks and provides a common framework for the potential design and control of regulatory systems in synthetic biology.

## Introduction

Regulation is considered one of a few defining features of living systems. Jacob and Monod (1961) described DNA as encoding the instructions necessary for the synthesis of individual proteins and additional “determinants” that promote or repress, i.e., regulate, protein synthesis. This duality of function was first articulated by Schrådinger in his influential book, What is Life, when he wrote, “They are law code and executive power – or to use another simile, they are architect’s plan and builder’s craft – in one”. The significance of regulation lies in its ability to control and execute a coordinated program of protein synthesis. Davidson and collaborators (Davidson and Erwin, 2006; Davidson, 2006) have advanced an explicitly computational perspective focusing on gene regulatory networks (GRNs) and developed a framework in which GRNs are represented in the form of wiring diagrams inspired by circuit diagrams in engineering. Modern systems biology adds a dynamical systems perspective to this static representation and large-scale detailed computer modeling (MacDonald et al, 1968; Churaev and Prokudina, 1989).

There is an extensive body of literature dealing with both the developmental and evolutionary history of regulatory networks and circuits in molecular biology, genetics, and metabolism. Nevertheless, widely shared features of regulation considered from the perspective of the corpus of essential regulatory elements have not been reviewed, and there is little consensus over a general model or specification of regulation that might span multiple levels of organismal organization.

The rare influential publications featuring “Theory of Regulation” in their title tend to focus on detailed immunological behavior in terms of immune network models that involve symmetrical stimulatory, inhibitory and killing interactions (Hoffmann, 1975). And outside of biology, the “Theory of Regulation” is common in the economics literature where it refers to the effect of regulatory policies in industries whose activities are strongly influenced by government intervention, see Armstrong and Sappington (2007) for a broad summary.

In many if not most genetic, proteomic, and epigenetic regulatory networks the most important feature of regulation is the control of rates of production and decay. Regulation therefore pertains to the direction or management of flux. An illuminating analogy is a system of traffic lights that regulates the flow of traffic by constraining and coordinating the rate of movement of traffic through a congested city. Traffic lights can be programmed to conform to a fixed schedule of delays that reflect historically observed local variations in traffic density. In the same way, genetic regulation can be assumed to reflect an evolved schedule of gene expression, and ensure an efficient response to a slowly changing environment through evolutionary adaptation. An analogous biological example would be a signaling cascade that transports an environmental signal (input) to the nucleus and converts this into transcriptional activity (output) in the face of numerous interfering reaction pathways. In practice, the cascade consists of many steps, each of which may be seen as a tiny regulatory system, ensuring efficient flow through the signaling network (Smith et al, 2011).

We adopt an approach in line with the objectives of synthetic biology which seeks to understand a system by building it. If we set out to build a regulatory system, what are the necessary and sufficient components to control the rate of gene and protein production in varying environments. In order to achieve this objective, and to ensure the maximum of clarity and simplicity, we focus on a single very well characterized regulatory system – the *lac* operon. We recognize that there are many clade-specific properties of gene regulation and seek to emphasize only those features that are shared across all kingdoms of life.

### Regulation Case Study: *lac* Operon

A carefully studied instance of a relatively simple regulatory system is the *lac* operon of *Escherichia coli*. The operon is something of a model system for regulation because it contains enough features to be representative of a larger class of regulatory mechanisms, and at the same time, simple enough to offer tractable insights into the basic elements of operation and their interactions. Moreover, the basic regulatory elements and their logic can often be generalized to encompass eukaryotic lineages.

The first insight derived from constructing a design specification language is that regulatory systems, even the same regulatory network, are often represented in widely different forms, and capture very different elements of regulation.

The *lac* operon in particular has been analyzed, and can be described as a metabolic regulatory circuit, using at least four different graphical representations (Fig. 1).

The first might be called the basic architecture of the operon (Fig. 1a). The architecture represents the essential components – genes and regulatory regions their locations, and some basic elements of their interactions and their dependencies. This representation is frequently observed in text books in genetics and general biology, *e.g.* in Krebs et al (2014).

The second is what is more commonly thought of as a regulatory network describing gene expression. (Fig. 1b). This representation includes fewer details than the architecture, does not indicate feedback, and seeks to capture the logic of regulation in terms of basic genetic interactions such as repression and induction. Regulatory networks sacrifice some degree of biological detail (such as the structure of the gene) in order to present simple insights into transcriptional regulation (Davidson and Erwin, 2006; Davidson, 2006).

A third representation is a causal influence diagram (Fig 1c). This does not represent direct physical interactions but causal interactions that could be mediated by additional components, not illustrated, indirectly. An advantage of representing the causal connectivity is that it conveys key features such as feedback but at the cost of remaining agnostic towards whether these interactions are positive or negative (Ay and Polani, 2008).

**Fig. 1.**
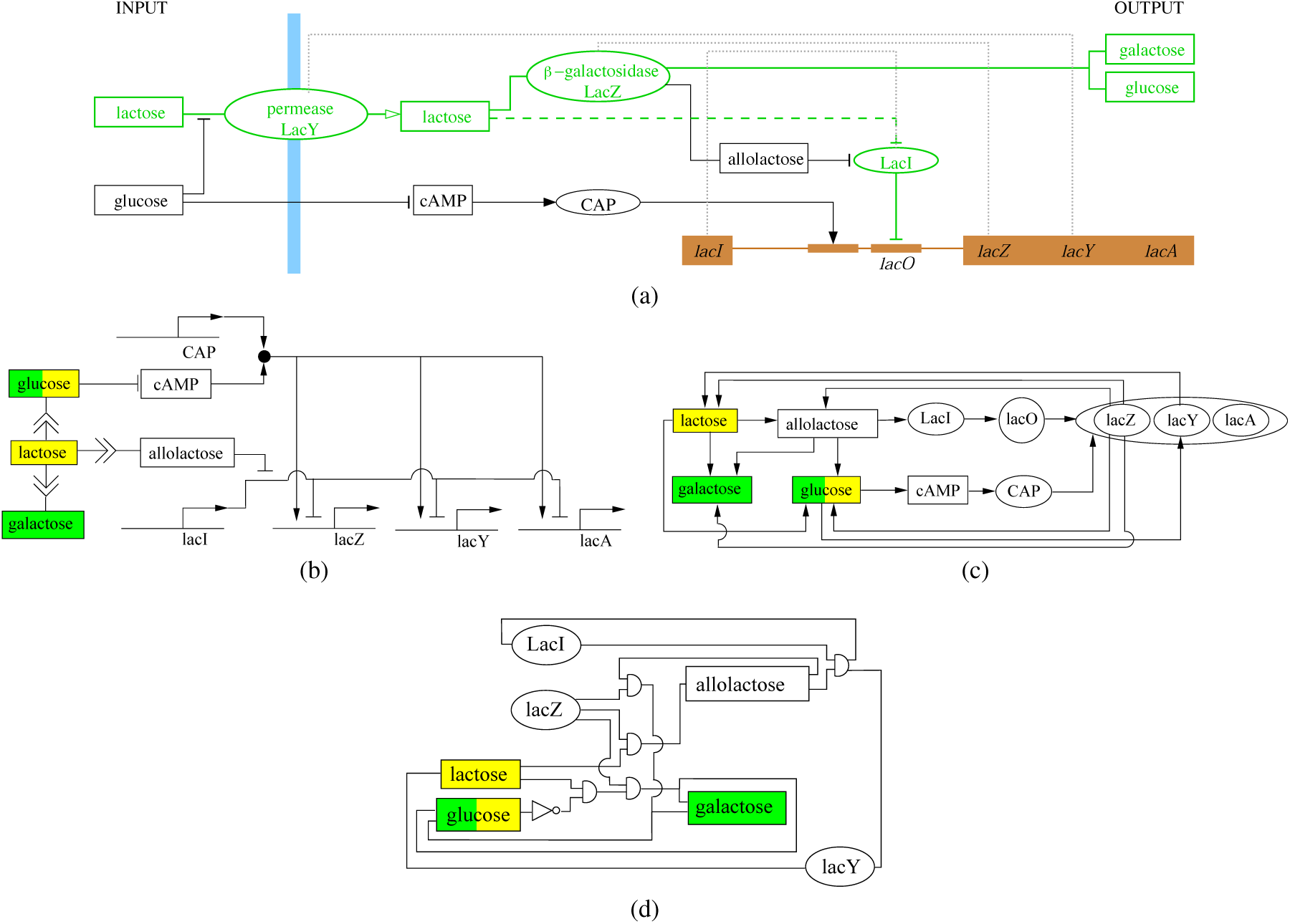
The *lac* operon through the lens of four common graphical representations. Lactose, taken up by the cell, is cleaved into glucose and galactose. (a) Basic Architectur of the lac Operon. In the classical representation (bold and green) the cell responds to the level of lactose by direct (dashed bold and green) interaction with LacI, the repressor, that shuts off operon expression in the presence of lactose through binding to lacO, the operator. A more complex picture emerges from the requirement of *beta*-galactosidase for the conversion of lactose into allolactose, and the involvement of glucose in the repression of cAMP-CAP activation of operon transcription (bold and black). Blue bar – cell membrane; brown – DNA sequence; wide brown boxes – protein conding regions; narrow brown boxes – cis-regulatory elements; ovals – proteins; rectangles – small molecules; grey dotted lines – gene expression; open arrowhead – tranlocation of lactose through the membrane by permease; full arrowhead – positive regulatory effect; flat arrowhead – negative regulatory effect; no arrowheads connect educts and products to enzymes. (b) **Regulatory Network according to Davidson et al.**Induction or repression of genes is indicated by arrows from regulatory molecules to the promoters of regulated gene. The outcome of any given combination of incoming signals is not represented. Only the effects on gene expression are illustrated and not the influences of gene expression on the abundance of small molecules. In this example, this generates a feedback-free diagram. (c) **Causal Influence Diagram** of the *lac* operon. An additional oval around the three gene products of the *lac* operon indicates that they are transcribed together (one mRNA, three independent open reading frames). Arrows indicate the influence of one element on another. It is impossible to determine whether influence is positive or negative. (d) **Logical Circuit Diagram** as deployed in synthetic biology. Logical relationship between molecules of the system are depicted using symbols from logical circuit diagrams in electronics. In the case of the *lac* operon only ’AND’ (half circle) and ’NOT’ (triangle) gates are required. In (b)-(d), inputs are marked in yellow while outputs are marked in green. Boxes represent small molecules, ovals represent proteins, and circles represent DNA elements. None of the three diagrams is able to capture dynamics.

A fourth representation takes the form of a formal logical network (Fig. 1d). This approach illustrates how inputs are combined, typically through Boolean functions such as AND and OR gates, to achieve a desired output. This logical, combinatorial insight is not available to any of the previous three representations as it moves beyond both simple induction, repression and causality, to include logic (Buchler et al, 2003; Silva-Rocha and de Lorenzo, 2008).

All of these representations largely (causal), or completely (architecture, GRN, logical) ignore dynamics. In order to capture the temporal features of regulation we need a different kind of illustration in which we explicitly capture the state of the system at successive points in time.

The dynamical representation (Fig. 2) is perhaps closest to the causal representation of regulation as it forces us to ignore details of architecture and logic for the sake of comprehensibility. There is no way of representing all of these essential elements using a single intelligible schematic. And this constraint of visualization frequently leads to one or more of the elements of regulation being neglected (West and Harrison, 2006).

**Fig. 2.**
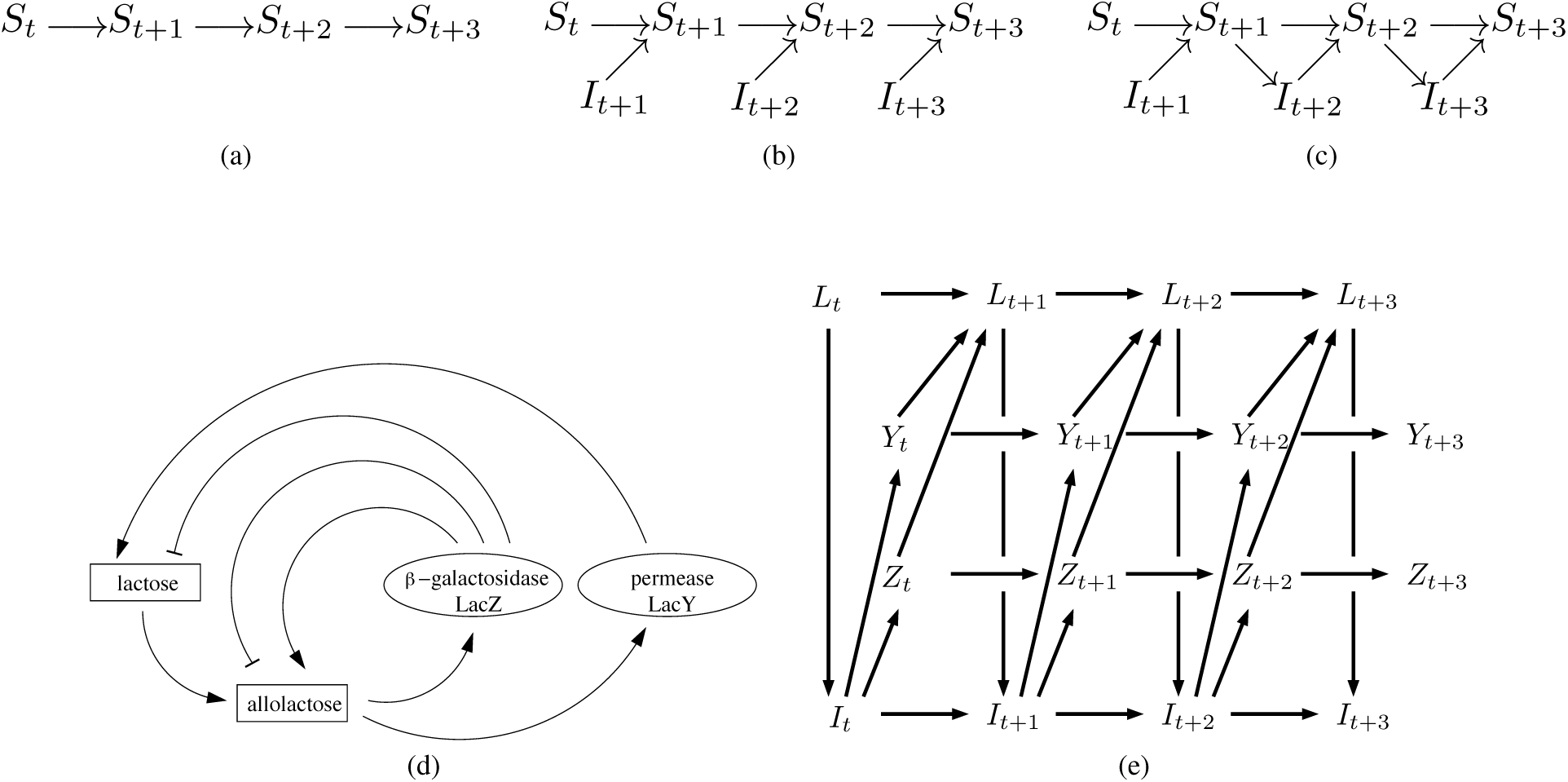
Boolean Networks: (a)-(c) illustrate the basic elements and architecture of a time expanded representation of a series of logical boolean mappings. Sequences of nodes with the same name represent temporal sequences (indicated by subscription) of the system’s state. Each arrow represents the ’flow’ of information between the states. (a) Internal state *St*+1 depends only on it’s previous state, i.e. *St*. (b) Internal state *St*+1 is influenced by it’s previous state and also by an input *I* from the environment at time *t* + 1. (c) As in (b) the system state *St*+1 is determined by the previous state *St* and the current input *It*+1. Additionally, the system state *St*+1 effects the input *I* at time *t* + 2 resulting in feedback. In this representation, feedback is shown as a directed path between temporally distinct nodes of the same state with at least on intermediate node in between. (d) & (e) Collapsed and time expanded boolean network for the *lac* operon. (d) The presence of a sufficiently high concentration of lactose drives expression of *β*-galactosidase LacZ and the permease LacY. Notice that *β*-galactosidase is an element of three feedback loops: lactose and allolactose metabolism (negative feedback) and transformation of lactose to allolactose (positive feedback). Expression of permease leads to an increase in intra-cellular lactose concentration by active transport into the cell. This in turn enhances permease expression (positive feedback). (e) Time expanded boolean network representation of the *lac* operon including only the feedback-associated elements. The lactose concentration (*L*) influences the amount of LacI (*I*) that can bind to lacO. As a function of bound LacI, expression of LacY (*Y*) and LacZ (*Z*) are enhanced or repressed. Expression of LacY leads to an increasing amount of lactose, whereas expression of LacZ catabolizes lactose. Note how complex the boolean network becomes even when restricted to the feedback associated elements in the mechanism. This explains, to some extent, why the dynamics are so frequently ignored.

By making explicit these five representations and their associated mechanistic implications, we are able through their union to highlight those elements and abstractions required to understand the underlying, time-varying and adaptive biochemistry of protein synthesis.

## Results

### Design Specifications for Regulation

None of these five representations is particularly effective when it comes to making comparisons among different regulatory systems. Even in conjunction they provide insufficient detail to reconstruct all that we know about any one biological system – such as the operon. They are in essence conceptual maps of regulation - they help us to navigate through complex biochemical pathways without being impeded by excessive detail.

In this paper we are proposing that there is value in a complementary approach – a design specification language for regulation. In engineering a design specification provides a detailed account of those factors, requirements and contexts necessary in order to manufacture an effective product. Design specifications provide a general language that enumerates critical components and their dependencies, which serve as lists of ingredients and recipes for construction. A design specification language helps us make progress towards a larger comparative framework. This framework would provide categorical terms for common elements allowing distinctions to be made between different regulatory networks in order to control or synthesize a regulatory system.

We introduce a regulatory design specification language by starting with the most elementary, effectively atomic, building block of regulation - the input-output function (illustrated in all five representations of Fig. 1), and adding properties as is required to capture more complex mechanisms. We culminate with a, “Typed IO dynamical system” which integrates elements of all five representations, and seeks to characterize rigorously the essential logical and dynamical elements and functions of any cellular regulatory mechanism. It is our objective to promote a common language and lexicon for regulation and not to provide a “final” definition which is likely to be of limited, and rapidly dated, value.

### I/O Maps

A very basic assumption is that all regulatory systems respond to inputs that are able to vary. These inputs can be external factors sensed by the system or internal states. Davidson’s gene regulatory network (GRN) framework of wiring diagrams assumes a form of effectively hardwired regulatory interactions (predetermined through evolution) that can be arranged into logical or Boolean building blocks. As aggregates, these can specify adaptive patterns (Bolouri et al, 2002; Materna and Davidson, 2007). In principle, variability can be continuous (a concentration or probability) or discrete. In its simplest form, the input is the absence or presence of a signal, typically a transcription factor. In response to the input a regulatory system provides a pre-determined output, typically a transcript. Such a system can conveniently be represented by an input/output map, Fig. 3(a).

**Fig. 3.**
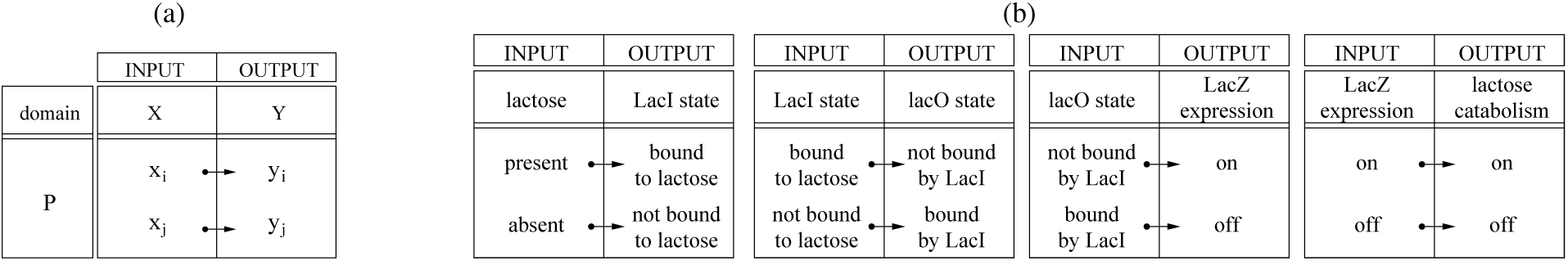
Typed I/O maps. (a) Formal representation. (b) A sequence of four I/O maps connects lactose availability (input of the first map) to lactose degradation (output in the last map). Notice that the input type of the successor is the output type of the predecessor. In the presence of lactose, LacI is bound by (a derivative of) lactose and dissociates from LacO. This allowes expression of *β*-galactosidase (LacZ) and, therefore, lactose catabolism. Notice also, that the first and the last I/O map are physically constrained while the maps two and three are “arbitrary” (see text for more detail.) In the example, discrete values are for simplicity.

An input-output (I/O) map is a mathematical or logical abstraction that relates the value of some observable *y* (output) to the value of some antecedent argument *x*(input). Traditionally this is described in terms of the domain *X*, the codomain *Y*, and *P* the set of ordered pairs that maps elements in *X* into elements in *Y*. In the context of regulatory systems, however, one needs to allow for stochastic I/O maps and a probability distribution of outputs as a function of an input *x*.

### Typed I/O Maps

The I/O map defines (a part of) any system’s dynamics by specifying how the value of *y* at a *subsequent* point in time depends on the *current* values of *x*. The I/O map therefore captures causal interactions. Domain *X* and co-domain *Y* can be taken as sets of values that are either numerical (expression levels, concentrations) or categorical (presence/absence) but cannot merely be identified with the real numbers or the integers. Instead, elements in X and Y are said to be of a particular type “X” and “Y”, respectively.

Formally, such types are simply sets that identify their elements by destinctive properties and define which operations can be performed on elements of the same set. A map can only take objects of a given type as its input and its outputs are also of a fixed type. For example, in mathematics, a variable can be typed as an integer, floating point number, or character. A more complex type is a function that can be “odd” or “even”, or a programming language that can be strongly or weakly typed. In all cases, typing provides for increased consistency and a reduction in ambiguity and error. All regulatory systems are dynamical systems that mediate the transmission of an input of one type to an output of another type.
Typed I/O maps can be concatenated provided the types match, i.e., *f*: *X → Y* and *g*: *Y → Z* can be combined to *f ◦g* : *X → Z* if and only if the co-domain of *f* coincides with the domain of *g*. This provides the formal foundation for decomposing systems into individual steps and for concatenating them into complex regulatory networks (Fig. 3(b)). In this way types provide us with a principled means of talking about levels in biology.

To summarize, the theory of typed I/O functions allows us to decompose a regulatory system into functional elements. The central dogma of molecular biology serves as a useful illustration. It represents two typed maps, DNA *→* RNA and RNA *→* protein, transcription and translation, respectively, that are concatenated at the level of RNA, to describe the process of gene expression.

For the case of the *lac* operon (Fig. 3(b)) one possible decomposition results in four maps where lactose abundance, transcripional repressor and operator states (LacI and lacO state), and levels of gene transcription and expression (lacZ expression) are the molecular types and the mappings reflect the molecular interactions.

How one goes about deriving meaningful types and maps is far from obvious. For example, consider the miRNA controlled degradation of mRNAs. Naively, we might simply classify the regulator (miRNA) as well as the regulated mRNA, as a single type – RNA. However, doing so we would not be able to derive an I/O map with different types for the input and output domain. In contrast destinguishing between miRNA, a type of RNA regulator, and mRNA, a type of regulated molecule, we can establish a mapping that describes miRNA target interactions. This emphasizes that it is not the species of polynucleotide that sets the type rather the operations permitted by the functional class of molecules.

### Encoding and Internal States

A key feature of many, but by no means all regulatory systems, is the presence of components that effectively act as internalized markers, or *encodings* of some adaptive aspects of the external environment. In the case of the *lac* operon (see Fig. 3(b)), the function of LacI is to sense the concentration of lactose and to provide this information in a form that is available as input for the *lac* operon. This tracking function is achieved through a conformational change of LacI upon binding of allolactose causing LacI to release the *lac* operator (lacO).

Internal encoding of external signals with markers or tracers serves several functions. The averaging of external fluctuations by an internal state (coarse graining) conveys increased robustness and resilience in the face of noise. More sophisticated encoding is able to filter out the detailed bio-chemistry of interactions in order to report only on bulk presence or absence, or classify interactions into distinct, categorical types. The functional value of a regulatory encoding is to allow for a separation of the machinery responsible for the sensory input from the subsequent decision making circuits – in other words, the creation of molecular classifiers.

In a formal sense, an encoding is a specialized type of I/O map that converts one type of object into another type of object. The purpose of the encoding is to render state information in a less noisy and more logical and hence universal, form. This is best known for the genetic code where RNA types are mapped into amino acid types with an associated error-correcting function.

Thus encoding promotes significant flexibility. Whereas a single type system, for example a metabolite concentration or temperature, is constrained largely by physical interactions, encodings among types promote considerable freedom for how information present in a physical signal can be processed, propagated, and aggregated towards down stream targets.

### Codes and Sets of Typed I/O Maps

#### Arbitrariness

A powerful feature of internal representations of the outside world is that there are few constraints on how features of the environmental information are mapped onto internal molecules and mechanisms. Moreover, typed I/O Maps are often “arbitrary” in the sense that there is a set of maps with the same I/O types that can produce different I/O mappings. This observation links typed I/O maps to the emerging field of code biology, where a “code” is viewed as “a set of rules that create a correspondence between two independent worlds” and that “are arbitrary in the sense that they are independent from physical necessity” (Barbieri, 2015). Within the framework of a design specification language, it is more natural to emphasize the map than the rules that specify a particular I/O map. This is because the very notion of arbitrariness becomes devoid of meaning in the context of a single mapping. Arbitrariness only makes sense for a code with a set of alternative I/O maps. The genetic code is the paradigmatic example. The genetic code can be rewired without loss of information by shuffling the assignment of amino acids to triplets, an exercise that has been carried out both in nature and in the lab (Chin, 2012; Neumann, 2012). Hence the genetic code is a code because it is one of a large family of possible realizations and not simply because it is an I/O map.

RNA regulators, such as miRNAs, recognize mRNA targets through complementary base pairing. This makes the code explicit and detaches in some form the mapping from the biochemical machinery (in this case Argonaute) mediating the interaction (Djuranovic et al, 2011; Mattick, 2004). This also allows for arbitrariness in the mapping, as the recognition sequences between mRNA and microRNA are in part the outcome of evolutionary contingencies. Likewise, the recognition of particular binding motifs by their corresponding transcription factors can be viewed as a code.

In the classical representation of the *lac* operon, lactose abundance induces lactose catabolism. The input-output relations can be represented by four maps, two of which are physically constrained and two are “arbitrary” (see Fig. 3(b)). Binding of lactose by LacI can only occur in its presence (Fig. 3(b), map 1) and lactose catabolism can only occure when LacZ is expressed (Fig. 3(b), map 4). These maps cannot be “rewired”. In contrast, the intermediate regulatory interactions constitute a system of negative inducibility (“repressing the repression”) that could theoretically be instantiated as a system “inducing the induction”. Instead of repressing LacZ expression through LacI *repressor* binding to the operator lacO in the absence of lactose, one could devise a LacI^*^ *inducer* that promotes LacZ expression upon binding to lacO^*^ in the presence of lactose. This is achieved by rewiring of I/O maps (Fig. 3(b), maps 2 and 3).

An extention of the *lac* operon representation includes glucose availability and the cAMP-CAP activation of operon transcription (see Fig. 1(a)). Concatenation of individual I/O maps results in a mapping of lactose/glucose presence as the input and operon expression as the output (see Fig. 4). It is not only an encoding of the external environment but a code because it maps a lactose/glucose absence/presence pattern to operon expression in a manner that is essentially arbitrary. The requirement that arbitrariness can only be derived from a set of maps with the same I/O types and different mappings and multifunctionality of regulatory types complicates the identification of true codes. A striking example is the “histone code”. Whereas the amount of histone acetylation marks, in general, promote DNA accessibility due to repulsive forces between negative charges on modified histones and the DNA backbone, particular histone acetylations seem to contribute to a more complex histone code Dion et al (2005). It is conceivable that some histone acetylation marks serve a dual role contributing to both, the “simple” (nonarbitrary) and complex histone code Prohaska et al (2010).

**Fig. 4.**
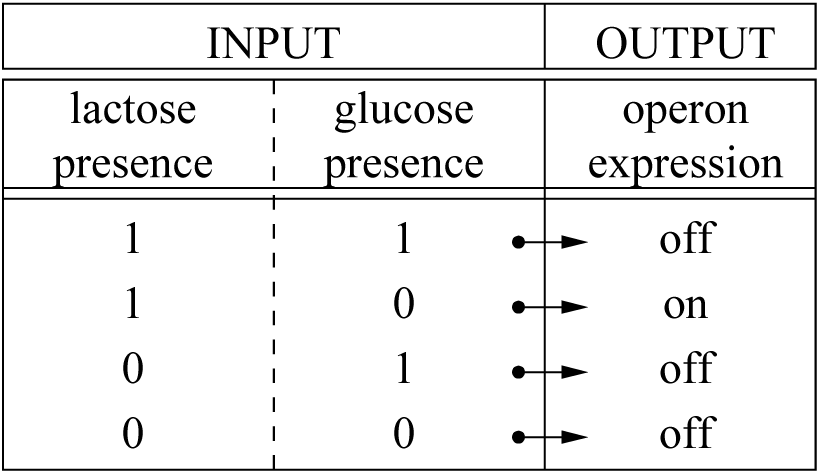
Code table for lactose and glucose availability and LacZ expression. Note that this mapping cannot be represented by any of the logical gates (i.e. AND, OR, XOR, NAND, NOR or XNOR) used in digital circuits.

Arbitrary mappings, and hence codes, are an essential element of synthetic biology which seeks to harness molecular machines to deliver alternative products.

A code need not be fixed, and the choice of I/O function can itself be sensitive to variations in the input domain. For example, modification of a transcription factor can modify expression of a set of regulated targets. The same applies to modification of tRNAs, which can change the binding affinity of the anti-codon for the codon. This form of arbitrariness is of key importance for the evolution of regulatory mechanisms. In system biology, the term “evolutionary plasticity” is often used for this type of arbitrariness of a code.

#### Combinatorial Architecture

Several distinct features contribute to arbitrariness and plasticity. A non-trivial combinatorial structure is often involved. In the genetic code the input are triples {*A,T,G,C*}^3^ (rather than just an unstructured set of 64 elements), leading to a different weighting for each of the three coding positions and the near complete redundancy of the third position. A combinatorially rich structure of the code-set provides multiple degrees of freedom in the mapping between input and output. This freedom in the typed I/O maps constitutes a target upon which evolution can act, modifying and diversifying the details of a regulatory system.

#### Indirection

A concept from computer science that can be useful in understanding typed I/O Maps, and biological codes in particular, is “indirection”. The key idea behind indirection is to decouple or reduce dependencies among processes. In computer science a good example of indirection is pointer arithmetic. A pointer is a value that encodes an address in memory. The pointer does not specify the contents of the memory only the location. Hence we might want to retrieve a value at a particular memory address stored by a previous process. This value might turn out to be a pointer to another address, and so forth. Function composition can be interpreted in the same manner: The pointer itself is an intermediate type *Y* which is set by a function *p* based on a input *X*. The function *q* resolves the pointer arithmetic such that the type *Y* is mapped onto the output of type *Z*. To regulate *Z* by *X*, for instance, the intermediate pointer type *Y* is used to mediate the link of *X* and *Z*.

Indirection in biological systems is observed in regulation of gene expression where regulation is achieved through binding of transcription factors to promoters or enhancers. Stressing the analogy to computer science, a transcription factor can be understood as a pointer that points to an address (binding site in the promoter) that contains gene expression values for a set of genes. For example transcription factors (TFs) that induce gene expression in specific sets of genes. Transcription factors, serving as the pointers, refer to addresses, i.e. promoters containing the transcription factor binding sites, that hold sets of genes as value.

Indirection decouples the selection of genes subject to expression change from the process of expression itself. An important effect of indirection is that changing pointer values changes the set of genes under regulation while adding new pointers puts the same set of genes under different control. Through this mechanism modification can affect large sets of genes in a coordinated fashion.

### Feedback

Perhaps the best known element of regulation, and the central dynamical component of typed I/O maps, is feedback. In the field of control theory and cybernetics, feedback is formally treated as a causal loop in which information about the past influences the behavior of a system into the future. Structurally, the output loops back into the input such that an event can “feed back” into itself (see Fig. 2d and 2e for feedback in the *lac* operon).

The key principle in the control of a thermostat, whose input-output properties can be set to arbitrary equilibrium temperatures, is feedback. The same is true of a traffic system that actively tracks vehicle density and uses this information to modulate stop-go intervals and thereby ensure flow.

In circuit diagrams, every directed cycle is a feedback loop. In temporally unfolded networks any directed path leading from *Xt* to *Xt*+*i* through intermediates signifies feedback. One can directly recover wiring diagram representations by simply identifying the nodes corresponding to the same component at different time points. Both, temporally unfolded networks and feedback loop diagrams for the *lac* operon are shown in Fig. 2.

Feedback loops have been studied extensively in biological networks. For example, mutual dependencies in large sets of co-regulated genes have been analyzed by sampling from an empirically supported distribution of coupling constants and connectivities. In this setting, the evolution of network architectures has been analyzed (Wagner, 1996; Bergman and Siegal, 2003), and small network motifs implementing particular types of feedback, such as feed-forward loops, have been identified (Alon, 2007).

Two important additional features of control are observability and controllability. A system is observable if from the output values of a system it is possible to determine the behavior of the entire system, including internal states. Thus for any possible sequence of state and control values, the current state can be determined in finite time using only the outputs. This is essential for the effectiveness of any selective process acting on regulation.

A system is controllable if there is a mechanism for moving a system around in its state or configuration space using a suitable controller. Complete controllability is achieved when an external input can move the internal state of a system from any initial state to any other final state in finite time. This is rather unlikely to be observed in nature. But some degree of controllability is required for adaptation to become possible.

Successful feedback requires both observability and controllability as the output needs to provide the key observable information to generate a feedback signal that can be used to control the future state of the system.

Feedback also introduces a complication into simple conceptual frameworks: the flow of causality is no longer linear. In particular, every node in a feedback loop regulates its successor and is regulated by its predecessor. The clear distinction of regulator and “the regulated” breaks down. This is perhaps the single most important limitation of static representations of regulatory principles.

### Memory

The fact that regulation is a dynamical system implies that there is at least a trivial form of memory present – namely the information in the current state that influences states at subsequent time steps (first order memory).

The phenomenon of hysteresis creates a more complex form of memory. Hysteresis describes a system in which an effect comes to depend not only on the current but also past inputs, i.e., some aspects of the system’s history. It is sometimes simplified as “an effect lagging behind the cause”. The salient point is that a full description of a system’s behavior requires not only the current input to be characterized but also information from the past. This information needs to be thought of as encoded in the internal state of the system as in a marker or tracer molecule.

More sophisticated mechanisms of regulation, or “regulated processes”, are not only responsive to a variety of forms of “external” stimulus, but are based on a cellular memory of past events. This is clearly the idea behind regulating an economy, where interest and exchange rates are modified in response to changing patterns of demand and trade, seeking to anticipate in advance probable changes to the patterns of individual behavior into the future.

Memory, as an element of regulation, also allows that a regulatory code is not stateless. A stateless code is one in which each I/O map is treated independently. Hence past regulatory events are not recorded in future acts of regulation. A stateful code with memory is one that is capable of adaptation or learning, and changes according to the elements it has experienced in the input domain, (Fig. 5).

**Fig. 5.**
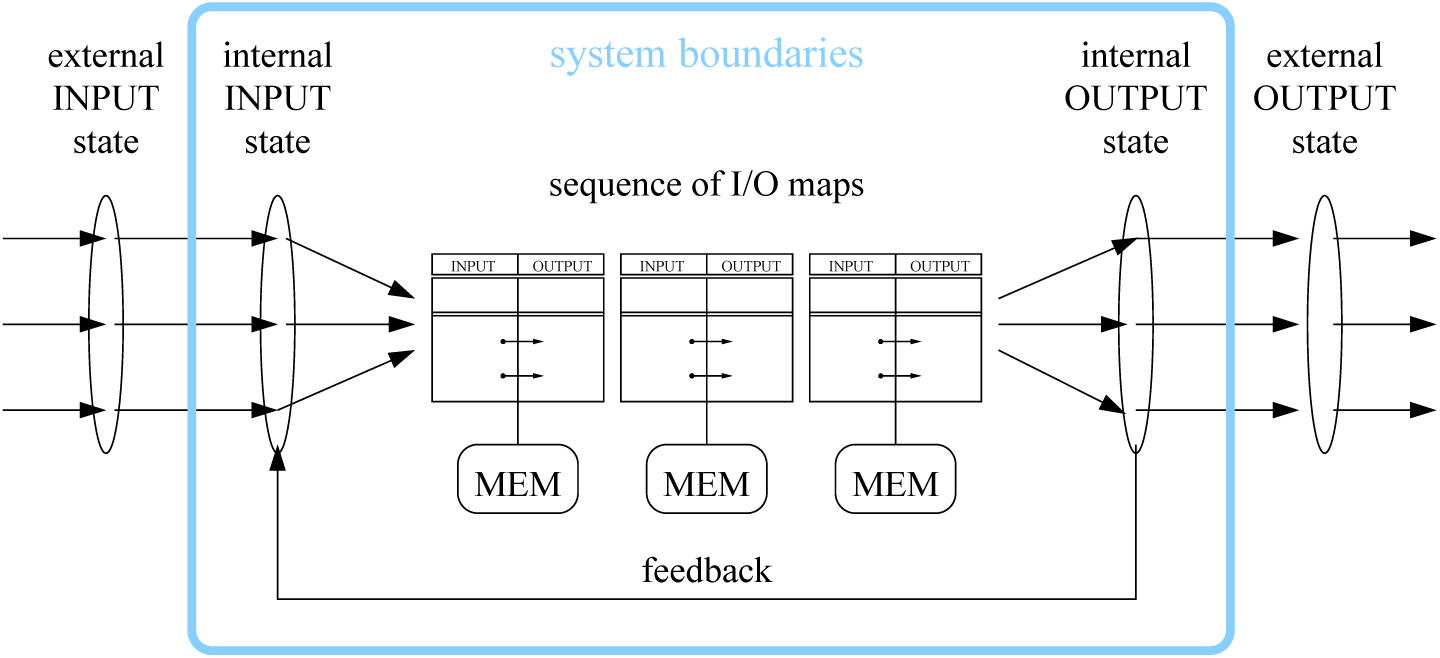
A regulatory system described in the proposed design specification language. From external input a regulatory system derives an internal representation that is than processed by the controler, a succession of I/O mappings, to generate an output. Feedback can link any intermediate output back to input of a predecessor as long as the types match. By copying processes, information can be written to and read from multiple independent memories (MEM).

A powerful memory device in the cell is the epigenome – the set of chemical modifications placed “on top of” the genome. Either directly attached to the DNA, as in DNA methylation, or to the histone proteins associated with DNA,e.g. histone acetylation, methylation, phosphorylation. These chromatin modifications can affect the expression of genes and restrict the transcriptional potential of a genome in a stable manner. Such effects have been observed at the transition from stem cells to differentiated cells (Chen and Dent, 2014). An emerging role of chromatin modifications is signal integration and storage (Badeaux and Shi, 2013). Mechanistically, the downstream ends of some internal and external signaling cascades results in the establishment of epigenetic marks. Those with no direct chemical effect on chromatin structure might still store the past presence of a signal. An even more complex form of memory makes use of feedback to ensure that an arbitrary I/O map remains stable in the presence of noise. This is memory in the sense of dynamical systems, in which the fixed points of a dynamical system are locally stable in the face of fluctuations in the density of some system variables.

Of great theoretical interest is the bistable switching of reaction networks based on a kinase and a phosphatase that can phosphorylate and dephosphorylate, respectively, a protein substrate at multiple sites. Rather than encode multiple variants of a single gene in a paralogous cassette at the level of the genome, only a single copy is encoded instead and the different states are determined by modifying the protein in a number of ways. Regulation is no longer described in terms of logic rules at the level of the DNA, but in terms of phase transitions following reconfiguration of enzyme reaction graphs (Krishna and Ramachandran, 1975). Prior history in such a system can be encoded and memorized as combinations of the distinct modification states. Similarly, the systematic use of chromatin modification likely serve as an explicit memory device (Prohaska et al, 2010; Arnold et al, 2013), i.e. the epigenetic memory, comparable to memory in present day computers Smith et al (2011).

## Discussion

The *lac* operon possesses all of the elements of a regulatory system outlined in the design specification. The operon has inputs that are mapped onto outputs. These mappings can be represented by sequences of typed I/O maps, some of which may represent codes. It is clear that the *lac* operon is a dynamical system sensing variation in the lactose and glucose concentrations in the environment as an input, controling lactose catabolism, and changing the lactose concentration in the environment as the output. Feedback loops are present, and through feedback the *lac* operon obtains a memory record of previous states of the environment.

Since all regulatory system can be analyzed in terms of typed I/O maps, this design specification makes it possible to compare very different regulatory mechanisms in which the elements of regulation are composed of distinct materials and components.

For example, histone acetylation regulates relaxation of condensed chromatin allowing for active transcription. An I/O map transforms the amount of histone acetylation on nucleosomes into transcription rates. When transcription takes place, histone acetylases are recruited that attach additional acetylation marks, generating positive feedback. There is as in the case of the *lac* operon, a stateless memory. At this level of abstraction emphasizing feedback loops, the *lac* operon appears to be very similar. A finer-grained representation serves to highlight the key differences. For example, there is a non-trivial code in the *lac* operon, whereas histone acetylations, in general, do not possess this feature.

The representation of regulatory systems in terms of typed I/O maps also behaves well under coarse graining or averaging. Changing the resolution with which the system is described leads to a transformation in the typed I/O maps by function composition. In the most coarse grained picture, the multiple feedback loops and typed I/O maps in the detailed model of the *lac* operon can be collapsed into one map (as in Fig. 4), still preserving the design elements of a regulatory system.

Regulation is a core organizing concept in biology. Regulation is described in a variety of different and complementary ways, from basic interaction rules, through to causal flows, logical networks, and dynamical systems. These are all useful approaches but they make comparison and construction of regulatory networks challenging. In our presentation we have emphasized the role of molecular (as well as more abstract) types and the relationships – maps – between them. This interpretation sets the stage for a complete formalization using the language of category theory (Spivak, 2014). We have argued in particular that our typed I/O maps allow controlled concatenations; hence they can be treated as maps (morphisms) in a formally precise category-theoretic sense. The prospect of a full formalization in this framework is not merely aesthetically pleasing. It promises the practical feasibility of a design specification language for regulation in terms of typed I/O maps. We have laid out a conceptual foundations for such a language. It consists in identifying the essential elements of regulation, and decomposing any given regulatory biochemistry into these elements. The later include, I/O functions, the explicit identification of types, the enumeration of codes, identification of memory mechanisms, and establishing the degree of observability and controllability, and hence the evolutionary plasticity of a target regulatory system.

## Acknowledgements

The authors thank the John Templeton Foundation for funding this research with the Grant “Origins and Evolution of Regulation in Biological Systems” ID: 24332. The opinions expressed in this publication are those of the author(s) and do not necessarily reflect the views of the John Templeton Foundation. This work was partially funded by the the German Federal Ministry of Science (0316165C) as part of the e:Bio initiative).

